## Supplementary figures and images for "The histone variant H2A.Z is required for efficient transcription-coupled NER and to maintain genome integrity in UV challenged yeast cells"

### Supporting Figure

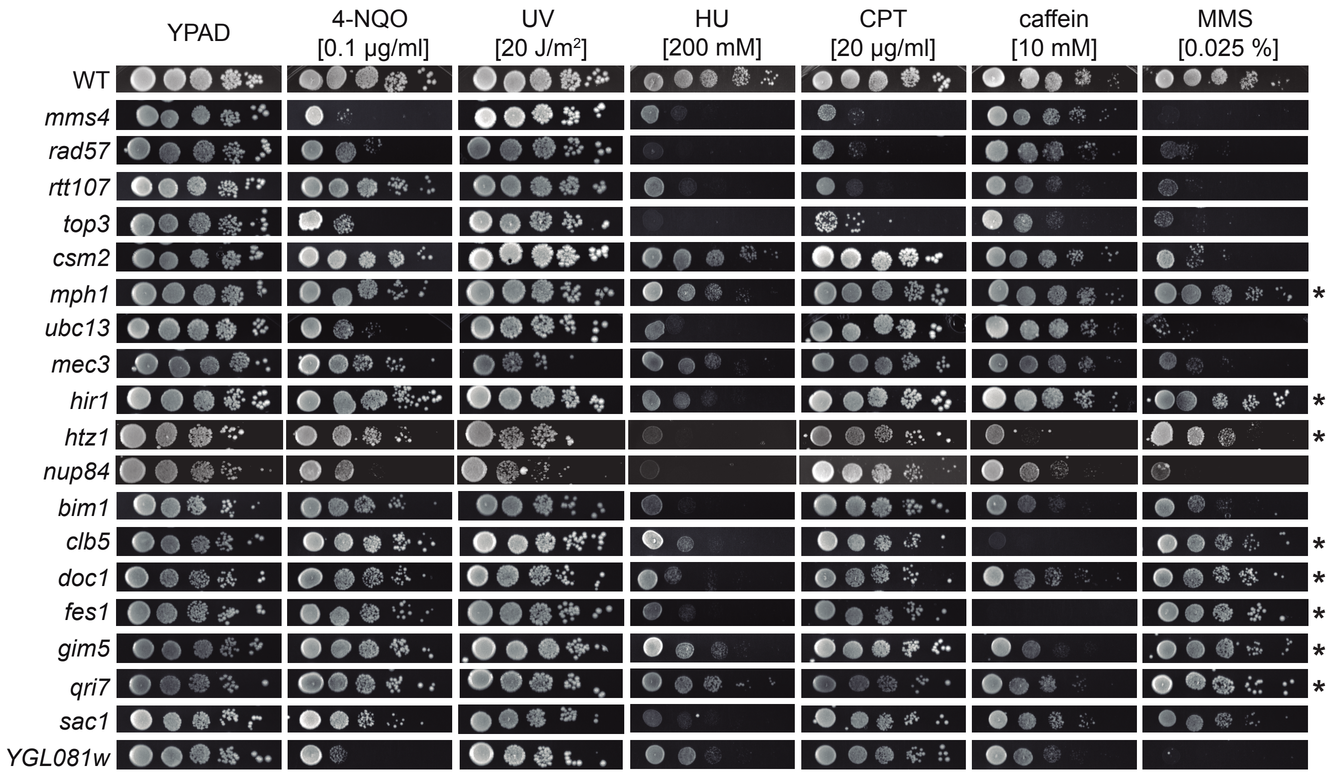


Supporting Figure S1

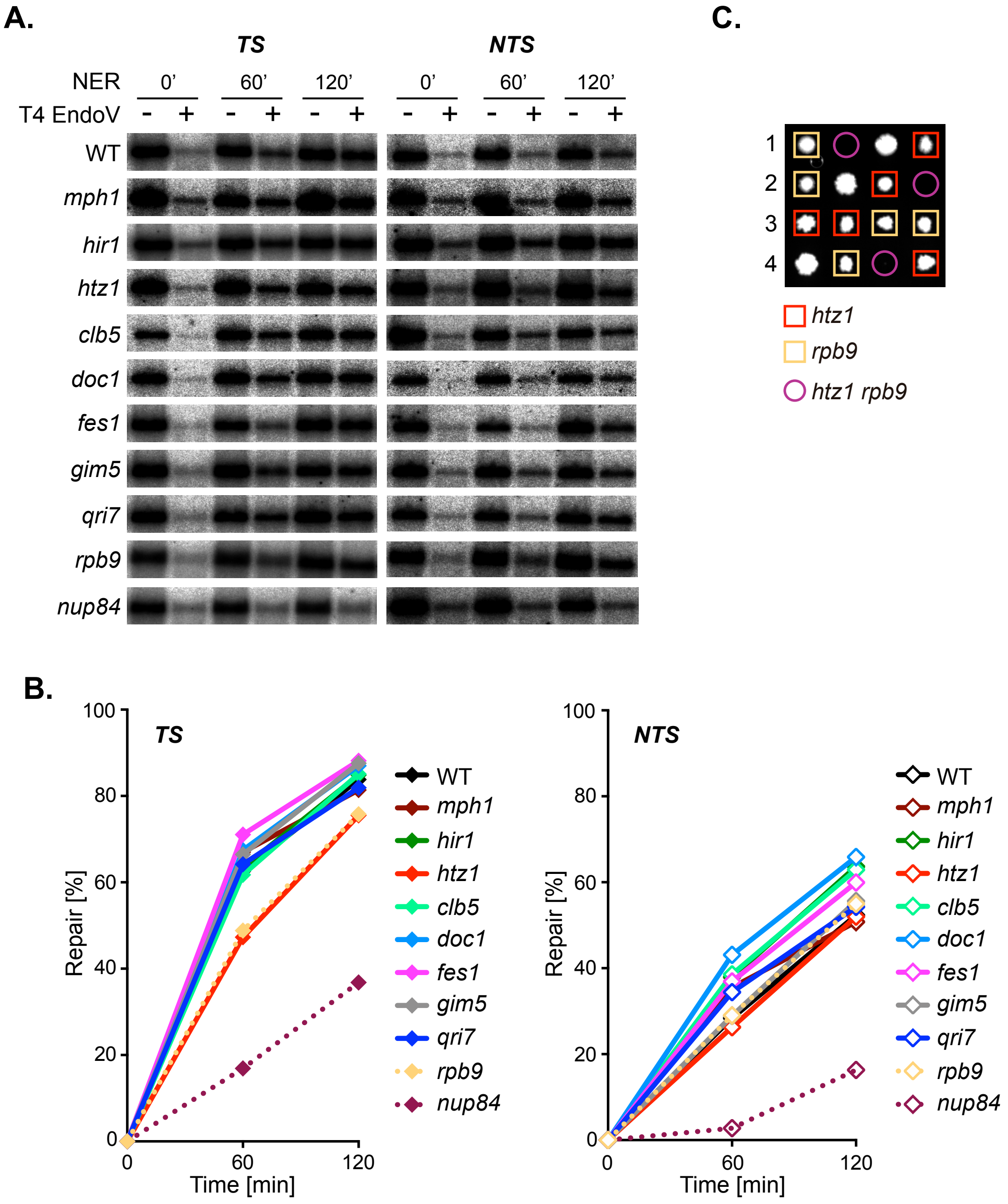


Supporting Figure S2

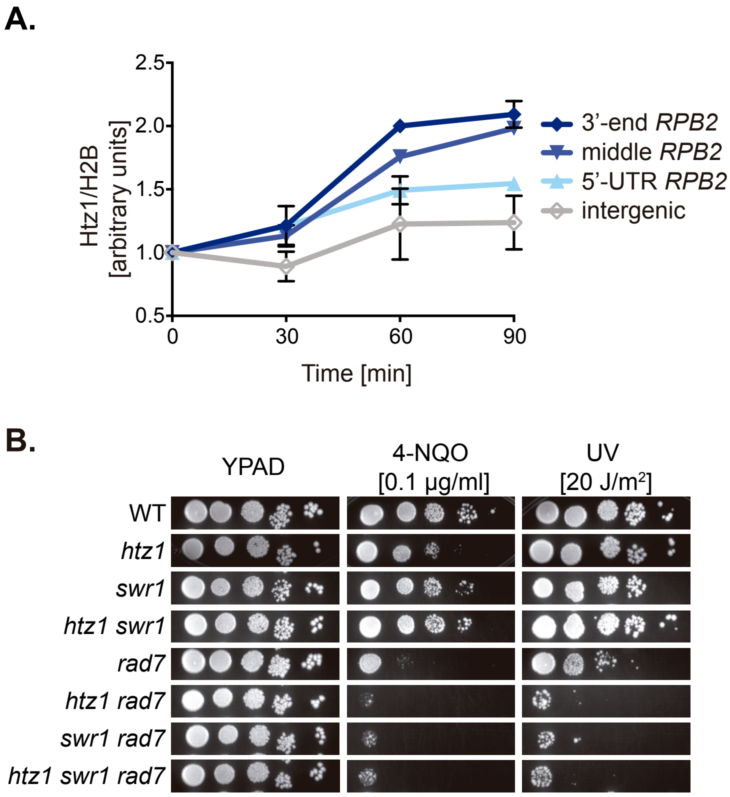


Supporting Figure S3

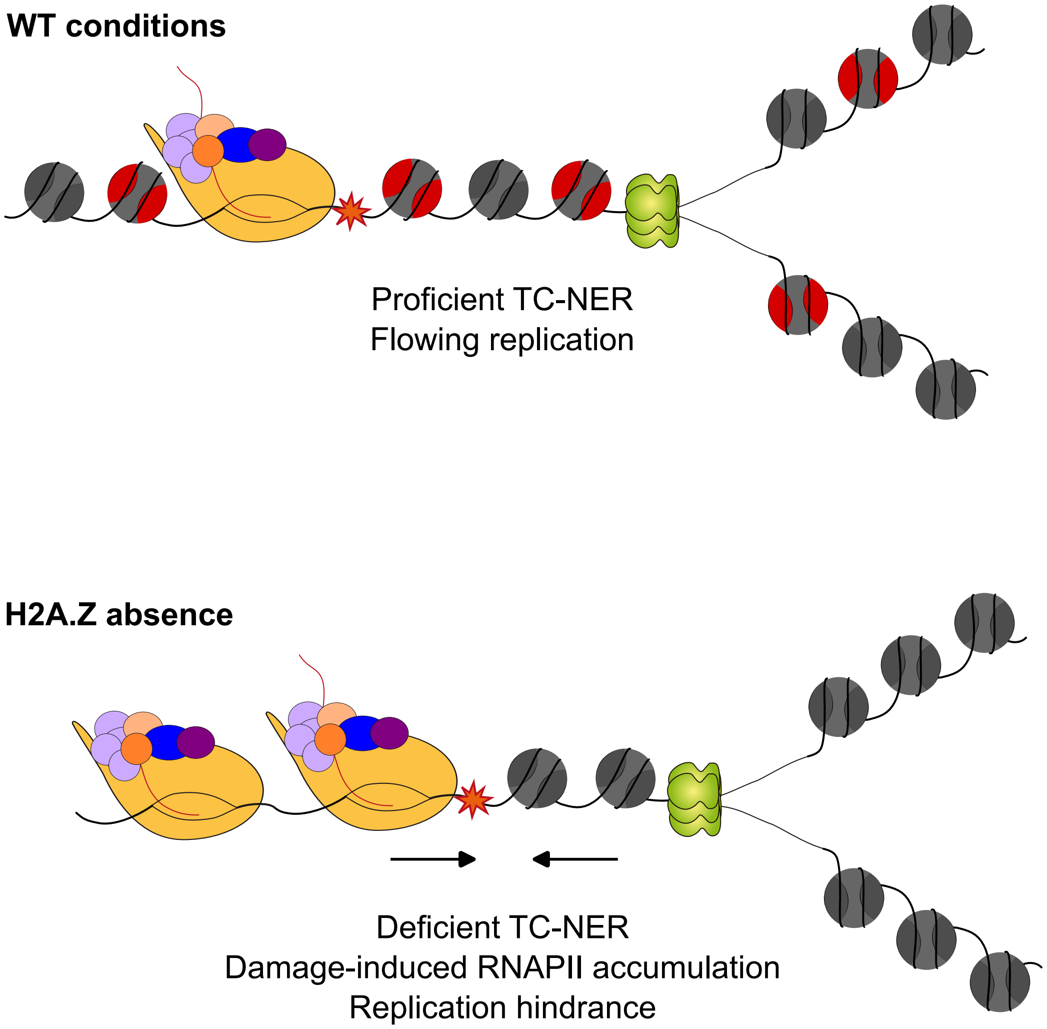


Supporting Figure S4
